# Neuromelanin accumulation drives endogenous synucleinopathy in non-human primates

**DOI:** 10.1101/2023.08.04.551615

**Authors:** Julia Chocarro, Alberto J. Rico, Goiaz Ariznabarreta, Elvira Roda, Adriana Honrubia, María Collantes, Iván Peñuelas, Alfonso Vázquez, Ana I. Rodríguez-Pérez, José L. Labandeira-García, Miquel Vila, José L. Lanciego

## Abstract

Although neuromelanin (NMel) is a dark pigment characteristic of dopaminergic neurons in the human substantia nigra pars compacta (SNpc), its potential role in the pathogenesis of Parkinson’s disease (PD) has often been neglected since most commonly used laboratory animals lack NMel. Here we took advantage of adeno-associated viral vectors encoding the human tyrosinase gene for triggering a time-dependent NMel accumulation within SNpc dopaminergic neurons in macaques up to similar levels of pigmentation as observed in elderly humans. Furthermore, NMel accumulation induced an endogenous synucleinopathy mimicking intracellular inclusions typically observed in PD together with a progressive degeneration of NMel-expressing dopaminergic neurons. Moreover, Lewy body-like intracellular inclusions were observed in cortical areas of the frontal lobe receiving dopaminergic innervation, supporting a circuit-specific anterograde spread of endogenous synucleinopathy by permissive trans-synaptic templating. In summary, the conducted strategy resulted in the development and characterization of a new macaque model of PD matching the known neuropathology of this disorder with unprecedented accuracy. Most importantly, evidence is provided showing that intracellular aggregation of endogenous alpha-synuclein is triggered by NMel accumulation, therefore any therapeutic approach intended to decrease NMel levels may provide appealing choices for the successful implementation of novel PD therapeutics.

## Introduction

The substantia nigra pars compacta (SNpc) is a pigmented structure macroscopically visible to the naked eye in human brain post-mortem samples. Pigmentation of dopaminergic neurons within the SNpc is driven by the age-dependent intracellular accumulation of neuromelanin (NMel), a dark-colored pigment resulting from a non-enzymatic auto-oxidation of dopamine^1,2^. Most importantly, a direct correlation between dopaminergic cell death and NMel content has long been established^3–5^. Since most commonly used laboratory mammalian animals such as rodents and rabbits lack NMel^6^, the potential role for NMel in the pathophysiology of Parkinson’s disease (PD) has often been neglected. In this regard, available clinical evidence showed a reciprocal association between patients with PD and melanoma, a cancer of pigmented skin cells known as melanocytes^7–19^. More importantly, detection of alpha-synuclein (α-Syn) in cultured melanoma cells and tissues derived from patients with melanoma has been recently reported elsewhere^20^.

The classical neuropathological scenario of PD is made of (i) a time-depending loss of pigmentation that correlates with neuronal loss^21^, (ii) presence of intracellular inclusions in pigmented neurons referred to either Lewy bodies (LBs; when located in the cytoplasm) or Marinesco bodies (MBs; intranuclear inclusions), and (iii) a pro-inflammatory scenario mediated by microglial cells. LBs are ring-shaped structures immunopositive for α-Syn and represent the main neuropathological hallmark of Parkinson’s disease. MBs are eosinophilic intranuclear inclusions considered as incidental findings observed in the brains of normal elderly subjects and are often observed in pigmented neurons in the SNpc^22,23^. Moreover, the so-called “Braak’s hypothesis” postulates a clinicopathological correlation of LB pathology and disease progression^24,25^ where pathology follows a predictable sequence of lesions from brainstem centers towards the cerebral cortex. Central to this hypothesis is the ability of α-Syn to spread transcellular through brain circuits, further inducing pathological aggregation of α-Syn in post-synaptic neurons by permissive trans-synaptic templating^26^.

Most of the currently available non-human primate (NHP) models of synucleinopathy are focused in recapitulating α-Syn aggregation by taking advantage of either adeno-associated viral vectors (AAVs) encoding different forms of the SNCA gene or using preformed α-Syn fibrils^27–32^. Although these models have important advantages when compared to traditional neurotoxin-based NHP models of Parkinson’s disease^33^, in particular when inducing a progressive dopaminergic neuronal degeneration triggered by α-Syn aggregation, α-Syn is mostly in the normal or disordered conformation and does not show morphological or biochemical features of Lewy body pathology. In this regard, an alternative approach has been recently introduced by taking advantage of AAVs encoding the human tyrosinase gene^34^ (hTyr). Upon AAV-hTyr delivery into the SNpc of rats, the enhanced expression of tyrosinase resulted in NMel pigmentation of dopaminergic neurons in the SNpc, an age-dependent PD phenotype, LB-like intracellular inclusions and progressive nigrostriatal degeneration. Since this model recapitulates all the neuropathological hallmarks typical of human PD with unprecedented levels of accuracy, here we sought to upgrade this rodent model to NHPs by following a similar strategy.

## Materials and methods

The collection of detailed protocols was deposited in protocols.io: https://www.protocols.io/private/0580290A56DB11EEB7B30A58A9FEAC02

### Study design

This study was aimed to develop and characterize a non-human primate (NHP) model of Parkinson’s disease mimicking the known neuropathological hallmarks of Parkinson’s disease to the best possible extent. Accordingly, we sought to determine whether AAV-mediated enhanced expression of human tyrosinase (hTyr) in the substantia nigra (SNpc) of non-human primates (NHPs) is able to induce a time-dependent accumulation of neuromelanin (NMel) in dopaminergic neurons, further triggering and endogenous synucleinopathy, progressive cell death and a pro-inflammatory scenario, in keeping with what was formerly reported in rats by taking advantage of a similar strategy^34^. Furthermore, the potential prionoid spread of endogenous alpha-synuclein (α-Syn) species towards the prefrontal cortex was analyzed, in an attempt to evaluate to what extent there is a propagation of endogenous α-Syn by permissive trans-synaptic templating (e.g. the so-called Braak hypothesis). Adult juvenile NHPs (*Macaca fascicularis*) were injected with adeno-associated viral vectors (AAVs) encoding either the hTyr gene (AAV-hTyr; delivered into the left SNpc) or a null construct for control purposes (AAV-null; injected into the right SNpc). In order to delineate a timeline for the underlying processes, one group of NHPs was sacrificed four months post-AAV deliveries (animals M308F4 and M310M4), whereby the follow-up timing for second experimental group was settled at eight months post-AAVs surgeries (animals M307F8 and M309M8). Neuroimage studies (MRI and MicroPET) were conducted *in vivo* at different time points. Upon animal sacrifices, brain tissue samples were processed for histological analysis comprising intracellular NMel levels, intracellular aggregates, nigrostriatal degeneration and neuroinflammation.

A graphical description of biochemical pathways underlying NMel synthesis is provided in Figure 1.

**Figure 1.**
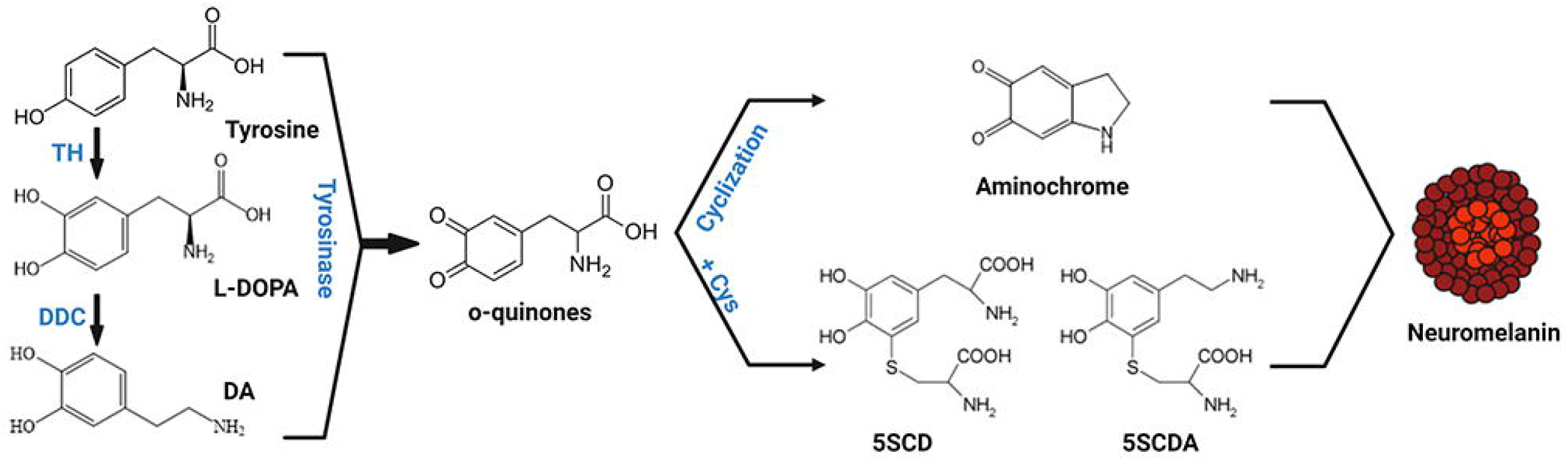
Schematic representation of neuromelanin synthesis. Dopamine is synthesized in cytoplasm from L-DOPA by DDC. Both cytosolica DA and L-DOPA can be oxidized either spontaneously or by tyrosinase to produce o-quinones. These, in turn generate aminochrome, 5SDCA and 5SCD which will act as precursors of the melanic components of neuromelanin. 5SCD, 5-S-cysteinyldopma; 5SCDA, 5-5cystenyldopamine; DA, dopamine; DDC, dopa-decarboxylase; L-DOPA, 3,4-dihydroxyphenylalanine; TH, tyrosine hydroxylase. Figure was created by Marta González-Sepúlveda with BioRender.com (agreement No. JI25JEO0Y7).

### Experimental animals

A total of four adult juvenile naïve *Macaca fascicularis* non-human primates (36-40 months-old; two males and two females; body weight 2.3-4.5 Kg) were used in this study. Animal handling was conducted in accordance to the European Council Directive 2010/63/UE as well as in keeping with Spanish legislation (RD53/2013). The experimental design was approved by the Ethical Committee for Animal Testing of the University of Navarra (ref: CEEA095/21) as well as by the Department of Animal Welfare of the Government of Navarra (ref: 222E/2021). Animals numbered as M307F8 and M308F4 are females, whereas animals M309M8 and M310M4 are both males. Animal records are shown in Supplementary Table 1.

### Viral vector production

Recombinant AAV vector serotype 2/1 expressing the human tyrosinase cDNA driven by the CMV promoter (AAV-hTyr) and the corresponding control empty vector (AAV-null) were produced at the Viral Vector Core Production Unit of the Autonomous University of Barcelona (UPV-UAB). In brief, AAVs were produced by triple transfection of 2 x 108 HEK293 cells with 250 μg of pAAV, 250 μg of pRepCap, and 500 μg of pXX6 plasmid mixed with polyethylenimine (Sigma-Aldrich). The UPV-UAB generated a pAAV plasmid containing the ITRs of the AAV2 genome, a multi-cloning site to facilitate cloning of expression cassettes, and ampicillin resistance gene for selection. Two days after transfection, cells were harvested by centrifugation, resuspended in 30 ml of 20 mM NaCl, 2 mM MgCl_2_, and 50 mM Tris-HCl (pH 8.5) and lysed by three freeze-thawing cycles. Cell lysate was clarified by centrifugation and the AAV particles were purified from the supernatant by iodixanol gradient as previously described^35^. Next, the clarified lysate was treated with 50U/ml of benzonase (Novagen; 1 h at 37 °C) and centrifuged. The vector-containing supernatant was collected and adjusted to 200 mM NaCl using a 5-M stock solution. To precipitate the virus from the clarified cell lysate, polyethylene glycol (Sigma-Aldrich) was added to a final concentration of 8% and the mixture was incubated (3 h, 4 °C) and centrifuged. AAV containing pellets were resuspended in 20 mM NaCl, 2 mM MgCl_2_ and 50 mM Tris-HCl (pH 8.5) and incubated for 48 h at 4 °C. The AAV titration method used was based on the quantitation of encapsulated DNA with the fluorescent dye PicoGreen®. Obtained vector concentrations were 1.7 x 10^13^ gc/ml for AAV-hTyr and 2.48 x 10^13^ for AAV-null. Plasmid map for pAAV-CMV-hTyr and sequence are provided in Supplementary Figures 1 & 2, respectively.

### Stereotaxic surgery for AAV deliveries

Surgical anesthesia was induced by intramuscular injection of ketamine (5 mg/Kg) and midazolam (5 mg/Kg). Local anesthesia was implemented just before surgery with a 10% solution of lidocaine. Analgesia was achieved with a single intramuscular injection of flunixin meglumine (Finadyne®, 5 mg/Kg) delivered at the end of the surgical procedure and repeated 24 and 48 h post-surgery. A similar schedule was conducted for antibiotic coverage (ampicillin, 0.5 mL/day). After surgery, animals were kept under constant monitoring in individual cages with ad libitum access to food and water. Once animals showed a complete post-surgical recovery (24 h), they were returned to the animal vivarium and housed in groups.

Stereotaxic coordinates for AAV deliveries into the SNpc were calculated from the atlas of Lanciego and Vázquez^36^. During surgery, target selection was assisted by ventriculography. Pressure deliveries of AAVs were made through a Hamilton® syringe in pulses of 1 μL/min for a total volume of 10 μL each into two sites in the SNpc, each deposit spaced 1 mm in the rostrocaudal direction to obtain the highest possible transduction extent of the SNpc. Once injections were completed, the needle was left in place for an additional time of 10 min before withdrawal to minimize AAV reflux through the injection tract. Coordinates for the more rostral deposits in the SNpc of AAV-hTyr (left SNpc) and AAV-null (right SNpc) were 7.5 mm caudal to the anterior commissure (ac), 5 mm ventral to the bicommissural plane (ac-pc plane) and 4 mm lateral to the midline, whereby the more caudal deposits were placed 8.5 mm caudal to ac, 5.5 mm ventral to the ac-pc plane and 4 mm lateral to the midline.

### Neuroimage studies

#### MicroPET Scans

MicroPET scans with (+)-α-[^11^C]Dihydrotetrabenazine (^11^C-DTBZ; a selective VMAT2 ligand) were performed on each animal at baseline and 1, 2, 4, 6 and 8 months post-AAV deliveries (6 and 8 time points only applying to animals with 8 months of follow-up). (+)-α-[^11^C]Dihydrotetrabenazine was synthesized by [^11^C]CH3 methylation of the corresponding (+)-9-O-desmehtyl-a-dihydrotetrabenazine precursor. [^11^C]methane was obtained by the wet chemistry method starting from [^11^C]CO2 porduced in the Cyclone 18/18 cyclotron at the Department of Nuclear Medicine, Clínica Universidad de Navarra, with a radiochemical purity of >95%. Images were acquired on a dedicated small animal Philips mosaic tomograph (Cleveland, OH, USA). The standard adquisition and quantification of the radiotracer binding potential was conducted as previously described^37^. In brief, a dynamic study of 40 min was acquired after the intravenous injection of the radiotracer. Obtained scans were analyzed by a radiotracer kinetic model using PMOD v3.2 software (PMOD Technologies, Ltd., Adliswil, Switzerland) to obtain parametric images containing the information of the binding potential of VMAT2. Parametric images were spatially normalized into standard stereotaxic space using a specific template^38^. The binding potential was measured using a predefined map of regions of interest (ROIs) defined over MRI images comprising the putamen nucleus. Changes in radiotracer binding potential were calculated for each animal at each time point.

#### MRI Scans

Studies were conducted at the Department of Radiology, Clínica Universidad de Navarra, in keeping with available protocols^39,40^. In brief, animals were scanned on a 3T MRI scanner (Siemens, Erlangen, Germany), using a 12-channel head array and consisted of the acquisition of an anatomical dataset and the NMel-sensitive sequence dataset with a total duration of 30 minutes. The anatomical T1-weighted image was acquired with MPRAGE sequence of 5 min duration. The following parameters were employed: 1 mm-isotropic resolution, FOV = 256 x 192 mm^2^, matrix = 256 x 192 voxels, 160 axial slices, repetition time/echo time = 1620/3.09 ms, inversion time = 659 ms, flip angle = 15°. Images of the SNpc were obtained with an NMel-sensitive T1-weighted fast spin-echo sequence with the following parameters: repetition time/echo time, 600/15 ms, two-echo train length, 11 slices, 2.0 mm slice thickness, 0.2 mm gap, 512 x 408 acquisition matrix, 220 x 175 field of view (pixel size 0.43 mm^2^, interpolated to 0.21 mm^2^), bandwidth 110 Hz/pixel, four averages, and a total scan time of 12 min. MRI studies were conducted four months post-AAV deliveries in all animals and eight months post-injection of viral vectors in animals M30F87 and M309M8.

#### Necropsy, tissue processing and data analysis

Anesthesia was firstly induced with an intramuscular injection of ketamine (10 mg/Kg), followed by a terminal overdose of sodium pentobarbital (200 mg/Kg) and perfused transcardially with an infusion pump. Animals M308 and M310 were sacrificed four months post-AAV deliveries, whereas animals M307 and M309 were euthanized eight months post-injection of AAVs. The perfusates consisted of a saline Ringer solution followed by 3,000 mL of a fixative solution made of 4% paraformaldehyde and 0.1% glutaraldehyde in 0.125 M phosphate buffer (PB) pH 7.4. Perfusion was continued with 1,000 mL of a cryoprotectant solution containing 10% glycerine and 1% dimethylsulphoxyde (DMSO) in 0.125 M PB pH 7.4. Once perfusion was completed, the skull was opened and the brain removed and stored for 48 h in a cryoprotectant solution containing 20% glycerin and 2% DMSO in 0.125 M PB pH 7.4. Next, frozen coronal sections (40 μm-thick) were obtained on a sliding microtome and collected in 0.125 M PB pH 7.4 as 10 series of adjacent sections. These series were used for (1) direct NMel visualization, (2) immunoperoxidase detection of TH, (3) dual immunofluorescent detection of tyrosine hydroxylase (TH) and P62 combined with brightfield visualization of NMel, (4) triple immunofluorescent detection of α-Syn, TH and P62 combined with brightfield visualization of NMel, (5) dual immunofluorescent detection of ubiquitin and P62 combined with brightfield visualization of NMel; (6) triple immunofluorescent detection of P62, Iba-1 and CD68 combined with brightfield visualization of NMel, (7) triple immunofluorescent detection of α-Syn (pre-digested with proteinase K), TH and P62 combined with brightfield visualization of NMel, (8) triple immunofluorescent detection of NeuN, TH and P62 and (9) multiple immunofluorescent detection of DAPI, α-Syn, P62 and TH.

A complete list of the used primary and bridge antisera (secondary antisera; either biotinylated or Alexa®-conjugated), together with incubation concentrations, incubation times and commercial sources is provided below:

##### List of reagents

| ITEM | DILUTION | INCUBATION TIME | SOURCE | IDENTIFIER |
| --- | --- | --- | --- | --- |
| **Antibodies** |  |  |  |  |
| Goat anti-Tyrosine Hydrox. (polyclonal) | 1:500 | Overnight | My Biosource | Cat# MBS421729 |
| Mouse anti-α-synuclein (monoclonal) | 1:40 | Overnight | Leica | Cat# NCL-L-ASYN |
| Mouse anti-P-α-syn (monoclonal) | 1:1000 | Overnight | Fujifilm Wako | Cat# 015-25191 |
| Guinea Pig anti-P62 (polyclonal) | 1:1000 | Overnight | PROGEN | Cat# GP62-C |
| Mouse anti-Ubiquitin (monoclonal) | 1:500 | Overnight | Abcam | Cat# ab7254 |
| Rabbit anti-NeuN (monoclonal) | 1:1000 | Overnight | Abcam | Cat# ab177487 |
| Rabbit anti-Iba 1 (polyclonal) | 1:500 | Overnight | Fujifilm Wako | Cat# 019-19741 |
| Mouse anti-CD68 (monoclonal) | 1:300 | Overnight | Dako | Cat# M0814 |
| Donkey anti-Goat (Biotin-SP) | 1:600 | 120 min | Jackson | Cat# 705-065-147 |
| Donkey anti-Goat (Alexa488) | 1:200 | 120 min | Invitrogen | Cat# A-11055 |
| Donkey anti-Goat (Alexa350) | 1:200 | 120 min | Invitrogen | Cat# A-21081 |
| Donkey anti-Goat (Alexa633) | 1:200 | 120 min | Invitrogen | Cat# A-21082 |
| Donkey anti-Guinea Pig (Alexa594) | 1:200 | 120 min | Jackson | Cat# 706-585-148 |
| Donkey anti-Mouse (Alexa488) | 1:200 | 120 min | Invitrogen | Cat# A-21202 |
| Donkey anti-Mouse (Alexa546) | 1:200 | 120 min | Invitrogen | Cat# A-10036 |
| Donkey anti-Rabbit (Alexa488) | 1:200 | 120 min | Invitrogen | Cat# A-21206 |
| **Commercial assays** |  |  |  |  |
| ABC kit standard |  | 60 min | Vector Labs | Cat# PK4000 |
| Neutral Red | 0.40% | 1 min | Sigma | Cat# 72210 |
| V-VIP |  | 1 min | Vector Labs | Cat# SK-4600 |
| Proteinase K | 1 μg/ml | 10 min | Invitrogen | Cat# 25530049 |
| DAPI | 1:50000 | 5 min | Invitrogen | Cat# D1306 |

#### Quantification of pigmented SNpc neurons

Every tenth section was counterstained with neutral red (NR) and used for estimating the number of transduced neurons with AAV-hTyr in the left SNpc. For this purpose, a deep-learning dedicated bi-layer algorithm was prepared with Aiforia® (www.aiforia.com), validated and further released (resulting in an error of 1.65% for quantifying either NMel+ / NR+ or NMel-/ NR+ neurons). Ten equally-spaced coronal sections covering the whole rostrocaudal extent of the SNpc were sampled per animal. Sections were counterstained with NR and scanned at 20x in an Aperio CS2 scanner (Leica, Wetzlar, Germany) and uploaded to Aiforia cloud. The boundaries of SNpc were outlined at low magnification (excluding neighboring areas such as the ventral tegmental area and the retrorubral field). The algorithm was then used as a template quantifying the desired neuronal populations. Representative images illustrating the accuracy of the conducted procedure are provided in Figure 3C1.

#### Quantification of NMel intracellular levels

Quantification of the intracellular density of NMel was achieved by measuring optical densitometry at the single-cell level with Fiji ImageJ software (NIH, USA) and converted to a logarithmic scale according to available protocol^41^. Scanned images comprising the entire rostrocaudal extent of the SNpc were inspected at a magnification of 80x. Analyses were conducted in 2,202 neurons for animal M307F8, 2,498 neurons in animal M308F4, 1,967 neurons in animal M309M8 and in 2,537 neurons for animal M310M4.

#### Assessment of nigrostriatal degeneration

The degree of nigrostriatal lesion resulting from NMel accumulation was measured both at origin and destination (e.g. at the level of the SNpc and striatum, respectively). At the level of the SNpc, every tenth section was stained for the immunoperoxidase detection of TH and used for estimating the number of number of TH+ neurons in the left and right SNpc. For this purpose, a deep-learning dedicated algorithm was prepared with Aiforia® (www.aiforia.com), validated and further released (resulting in an error of 4.82% for quantifying TH+ neurons). Ten equally-spaced coronal sections covering the whole rostrocaudal extent of the left and right SNpc were sampled per animal. Sections were scanned at 20x in a slide scanner (Aperio CS2; Leica), uploaded to Aiforia cloud and TH+ neurons were quantified according to a similar procedure described above for estimating the number of NMel+ neurons. Regarding nigrostriatal degeneration at destination, up to 25 equally-spaced coronal sections stained for TH covering the whole extent of the left and right caudate and putamen nuclei (pre- and post-commissural putamen and caudate) in each animal were scanned at 20x and used for measuring TH optical densities with Fiji Image J and converted to a logarithmic scale.

#### Analysis of NMel intracellular density and neuronal inclusions

In order to evaluate to what extent intracellular NMel levels correlate to the presence of intracellular inclusions, sections comprising all SNpc levels and stained for TH and P62 were used. The immunofluorescent detection of TH and P62 was combined with brightfield visualization of neuromelanized neurons under the confocal microscope at a magnification of 63x. Intracellular NMel levels were measured as described above by taking advantage of confocal Z stacks of similar thickness. For every single animal, a minimum of 25 pigmented dopaminergic neurons showing P62 aggregates were randomly selected from each section (comprising 12 equally-spaced sections per animal covering the whole rostrocaudal extent of the SNpc) and compared with a similar number of NMel+ / TH+ neurons without P62 intracellular inclusions.

#### Quantification and statistical analysis

Statistical analysis was done in GraphPad Prism version 9.0.2. for Windows and Stata 15 (Stata Corp. 2017. Stata Statistical Software Release 15, College Station, TX; StataCorp LLC). Relevant tests are listed in figure legends. Species with *p* < 0.05 were considered statistically significant.

### Data availability

Further information and request for resources and reagents should be directed to and will be fulfilled by the corresponding author, Jose L. Lanciego. Full datasets can be found at Zenodo repository (DOI: 10.5281/zenodo.8359416).

## Results

### Neuroimage studies

MicroPET scans with (+)-α-[^11^C]Dihydrotetrabenazine (^11^C-DTBZ; a selective VMAT2 radioligand) were performed on each animal at baseline and 1, 2, 4, 6 and 8 months post-AAV deliveries (6 and 8 months time points only applying to animals with 8 months of follow-up). For each animal and time point, the binding potential of ^11^C-DTBZ (reflecting nigrostriatal VMAT2+ innervation) was measured in MRI-matched pre-defined regions of interest across 10 consecutive sections comprising the whole rostrocaudal extent of putamen nucleus. Upon deliveries of AAV.hTyr into the left SNpc, significant declines in radiotracer binding potential were observed in the putamen nucleus from both female animals (animals M308F4 and M307F8). By contrast, measurements performed in male animals (animals M310M4 and M309M8) most often failed to reach statistical significance (Figure 2A).

**Figure 2.**
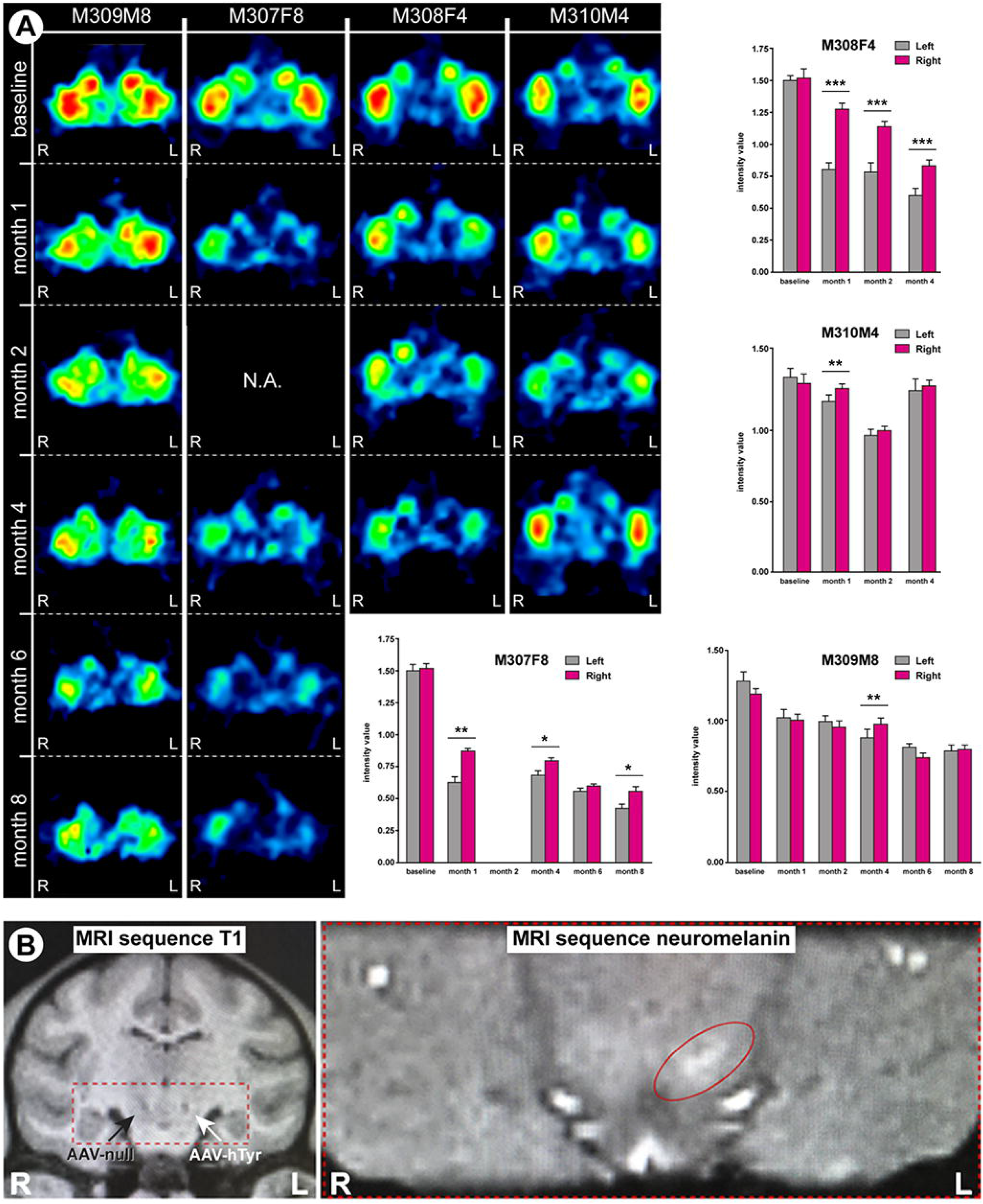
Neuroimage studies. (A) MicroPET studies. Representative coronal sections of the NHP brain at the level of the putamen nucleus showing the binding potential of ^11^C-DTBZ (a selective VMAT2 ligand). Histograms illustrate the obtained measurements by comparing radiotracer uptake in left vs. right hemispheres at each pre-defined time points. Statistical differences were more often found in female animals. \**p* < 0.05, \*\**p* < 0.01, \*\*\**p* < 0.001, related samples *t* test, n = 10 sections/animal. Data are represented as mean +/-SD. (B) MRI scans. Left: Anatomical T1-weigthed coronal MRI scan taken at the level of the ventral mesencephalon showing the location of the injection sites for AAV-hTyr (left SNpc) and the control AAV-null (right SNpc). Right: Hyperintense area in the left SNpc as observed with a neuromelanin-dedicated sequence.

Anatomical T1-weighted MRI images were acquired to further evaluate the accuracy of the viral vector deliveries, comprising AAV-hTyr and AAV-null deposits into the left and right SNpc, respectively. Next, animals were scanned with a NMel-dedicated sequence enabling the visualization of a hyperintense area at the level of the left SNpc (Figure 2B), this hyperintense area indicating NMel accumulation.

### Neuromelanin accumulation induced by hTyr overexpression

In all animals, the AAV-mediated enhanced expression of hTyr resulted in a macroscopically-visible pigmentation of the left SNpc that can be observed as a darkened area throughout the whole rostrocaudal extent of the left SNpc. By contrast, the delivery of the control AAV into the right SNpc resulted in a complete lack of pigmentation. Moreover, when comparing animals with a follow-up time of 4 vs. 8 months, a time-dependent loss of pigmentation was observed, more evident in female animals (Figure 3A).

**Figure 3.**
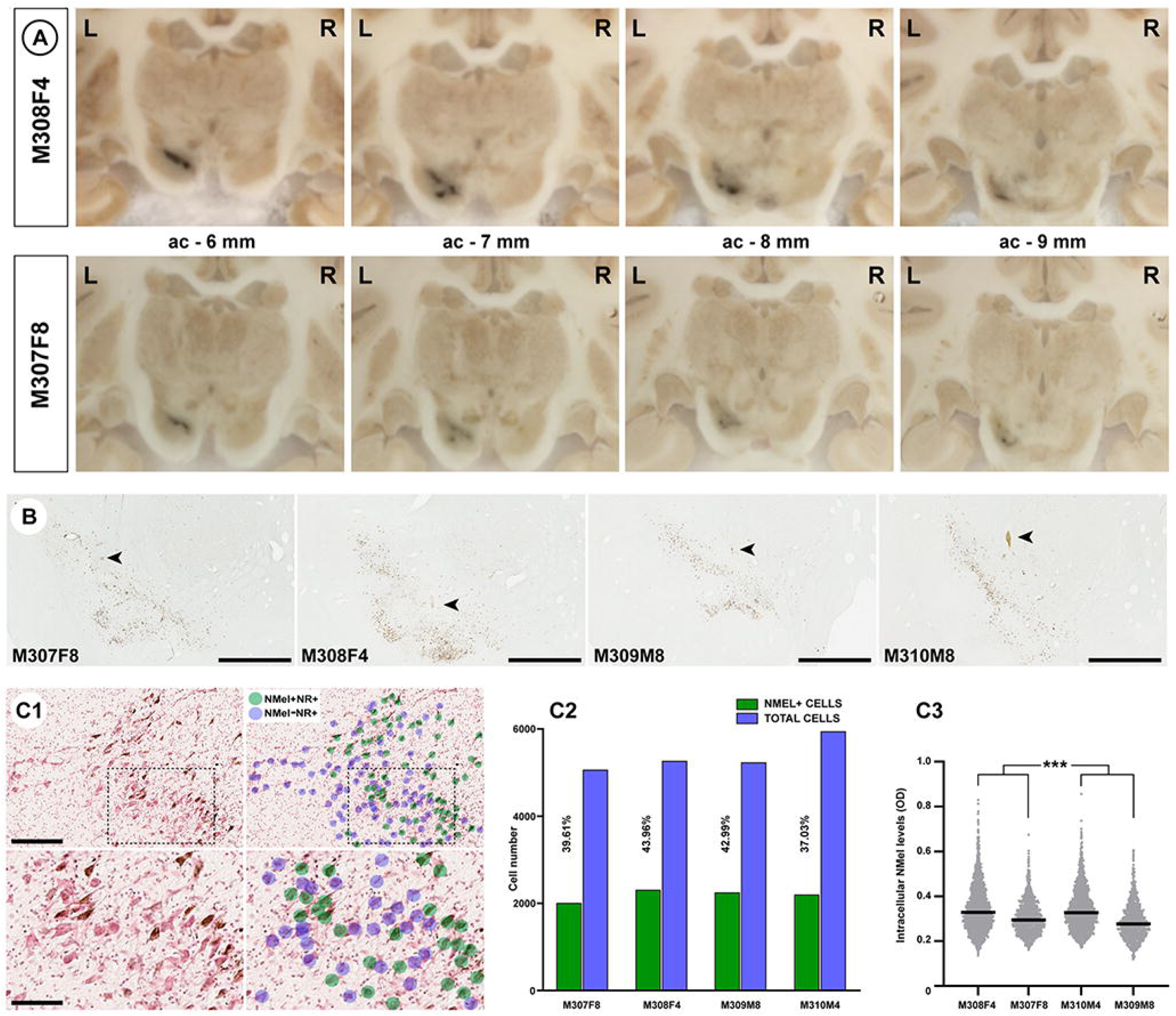
Pigmentation of the substantia nigra upon the delivery of AAV-hTyr. (A) Macroscopic images of NHP brains taken from the microtome during sectioning. A darkened area at the level of the left SNpc is visible to the naked eye throughout the whole rostrocaudal extent of this structure. A time-dependent loss of pigmentation is observed when comparing animals with post-viral injections follow-up times of 4 and 8 months. (B) Low-power microphotographs taken from non-stained sections of all animals at the level of the SNpc. Pigmentation with NMel is clearly visible even at low magnification. Arrowheads indicate the location of the injection sites. Scale bar, 1,000 μm. (C1) Illustrative example showing the conducted procedure for quantifying the number of neurons being transduced with AAV-hTyr in the SNpc. By taking advantage of sections counterstained with neutral red (NR), a bi-layered algorithm was prepared with Aiforia® (www.aiforia.com) to further disclose pigmented and non-pigmented NR+ neurons in the SNpc. (C2) Histogram showing the percentages of pigmented neurons in the SNpc resulting from the use of the Aiforia® algorithm. (C3) Boxplot showing the distribution and mean values of intracellular NMel levels in all animals. NMel densities in female animals are higher than in males at each time-point post-injection of AAV-hTyr. \*\*\**p* < 0.001, nested ANOVA test with time & gender as fixed factors, and monkeys nested within fixed factors. Measurements were conducted in 12 consecutive sections covering the entire rostrocaudal extent of the SNpc (equally spaced 400 μm each).

Microscopical examination of tissue samples at the level of the ventral mesencephalon revealed NMel intracellular accumulation in neurons of the SNpc, this accumulation reaching levels high enough to avoid the use of histochemical stains enhancing NMel signals such as the Masson-Fontana method (Figure 3B). The percentage of SNpc cells transduced with AAV-hTyr was estimated by taking advantage of a dedicated bi-layered algorithm disclosing pigmented vs. non-pigmented neurons in sections counterstained with neutral red (Figure 3C1). On average, 40% of cells in the SNpc displayed NMel accumulation, ranging from 37.03% in animal M310M4 (male, 4 months of follow-up) to 43.96% in animal M308F4 (female, 4 months of follow-up), these values likely reflecting a roughly similar accuracy for the conducted stereotaxic deliveries of viral vectors (Figure 3B). That said, it is also worth considering that the obtained percentages of cellular transduction might be somehow underestimated, bearing in mind that NMel accumulation cannot longer be measured in degenerated neurons, in particular for animals with 8 months of post-injection follow-up periods (M307F8 and M309M8). Moreover, a variable cell-to-cell degree of pigmentation was observed. Accordingly, the level of intracellular NMel accumulation was estimated at the single-cell level in all animals across 12 consecutive sections (equally spaced 400 μm) covering the entire rostrocaudal extent of the left SNpc. Obtained data showed that female animals exhibited a higher mean level of intracellular NMel than male subjects at both 4 and 8 months of follow-up, with a slight decline in animals with longer follow-up times (M307F8 and M309M8; Figure 3C3).

### Nigrostriatal neurodegeneration

To further address to what extent NMel accumulation is leading to dopaminergic cell neurodegeneration, tyrosine hydroxylase-positive neurons (TH+) in the left and right SNpc were quantified with a dedicated Aiforia® algorithm. Loss of TH+ dopaminergic cells was noticed in both female animals, ranging from 47.13% cell loss in animal M308F4 to 49.61% in animal M307F8, whereas much lower levels of degeneration were observed in male animals (8.63% of reduction in M310M4 and 19.93% in M309M8). In female animals, loss of TH+ cells was maintained throughout the entire rostrocaudal extent of the left SNpc (Figure 4A). As expected, the observed dopaminergic cell loss in the SNpc is associated to a reduction in TH+ terminals in the caudate and putamen nuclei, as measured with optical densitometry in 20 consecutive coronal sections (equally spaced 400 μm) covering all the pre- and post-commissural striatal territories in both hemispheres. A significant loss of TH+ terminals was found at the level of the caudate and putamen nuclei in animals M307F8 and M308F4. For both nuclei, higher differences were constantly found in post-commissural locations (Figure 4B). Optical densities of TH+ terminals in the two male animals failed to reach statistical significance at the level of the caudate and putamen nuclei, without any apparent difference between either pre- or post-commissural locations.

**Figure 4.**
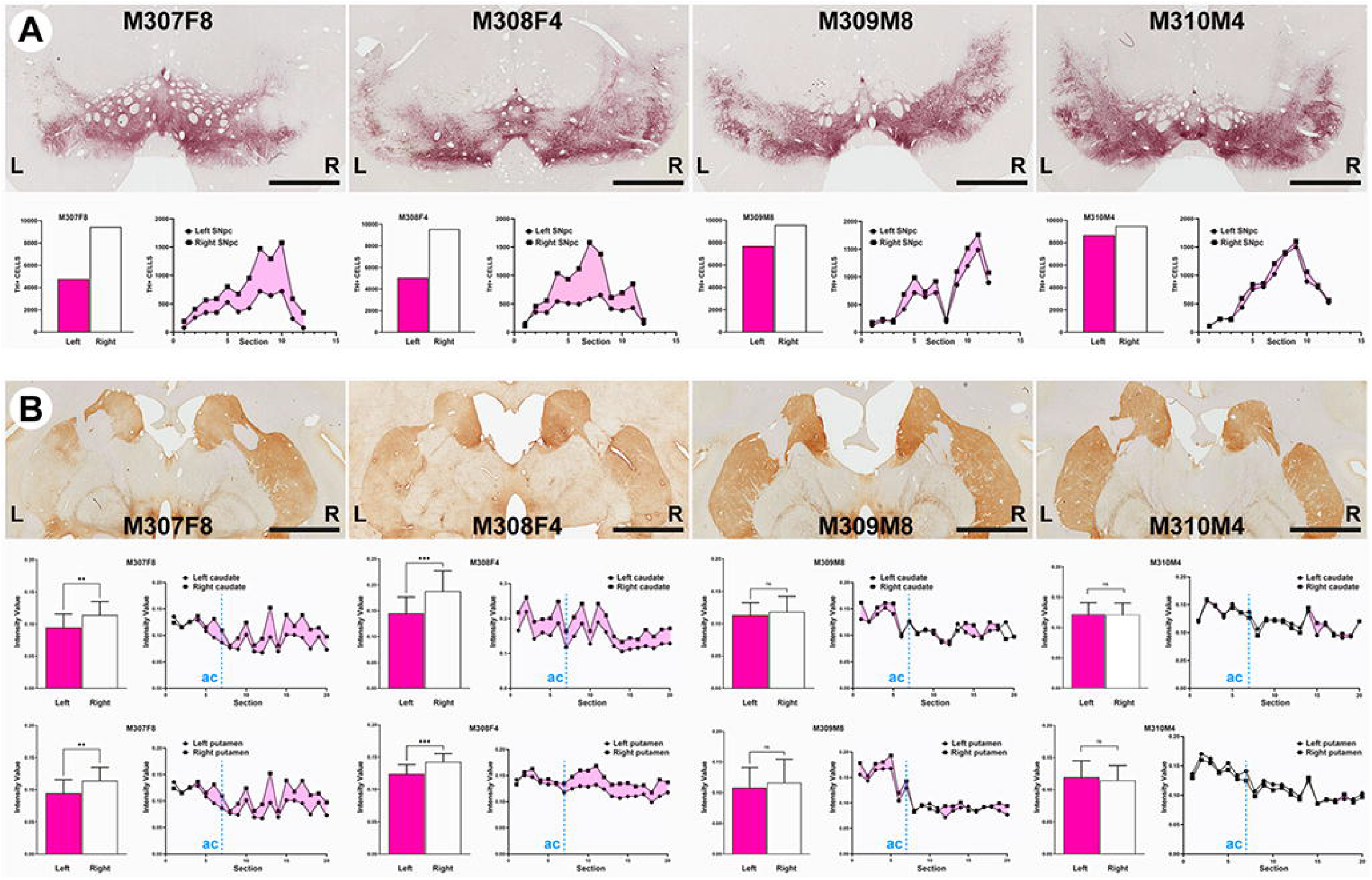
Time-dependent dopaminergic nigrostriatal damage driven by neuromelanin accumulation. (A) Coronal sections taken at the level of the SNpc immunostained with an antibody against tyrosine hydroxylase (TH) and visualized with a purple peroxidase chromogen (V-VIP). Histograms below illustrate the number of TH+ cells in the left vs. right SNpc, together with the rostrocaudal distribution of cell numbers in the SNpc (www.aiforia.com). Scale bar, 3,200 μm. (B) Coronal sections taken at the level of the post-commissural caudate and putamen nuclei immunostained for TH and visualized with a brown peroxidase chromogen (DAB). Scale bar, 6,400 μm. Histograms below illustrate optical densities of TH in the left and right caudate and putamen nuclei for each animal, together with the rostrocaudal distribution. Dashed blue lines labeled as ‘ac’ indicate the position of the anterior commissure. \*\**p* < 0.01, \*\*\**p* < 0.001, unpaired *t* test, n = 20 sections/animal. Data are represented as mean +/-SD.

Upon degeneration of pigmented dopaminergic neurons, NMel is released to the extracellular space as observed in the left SNpc in all animals. In keeping with similar phenomena observed in postmortem PD samples^44^, extracellular NMel deposits were often found in perivascular locations (Figure 5A). Extracellular NMel granules were found inside phagocytic cells positive for Iba-1 and CD68 (Figure 5B).

**Figure 5.**
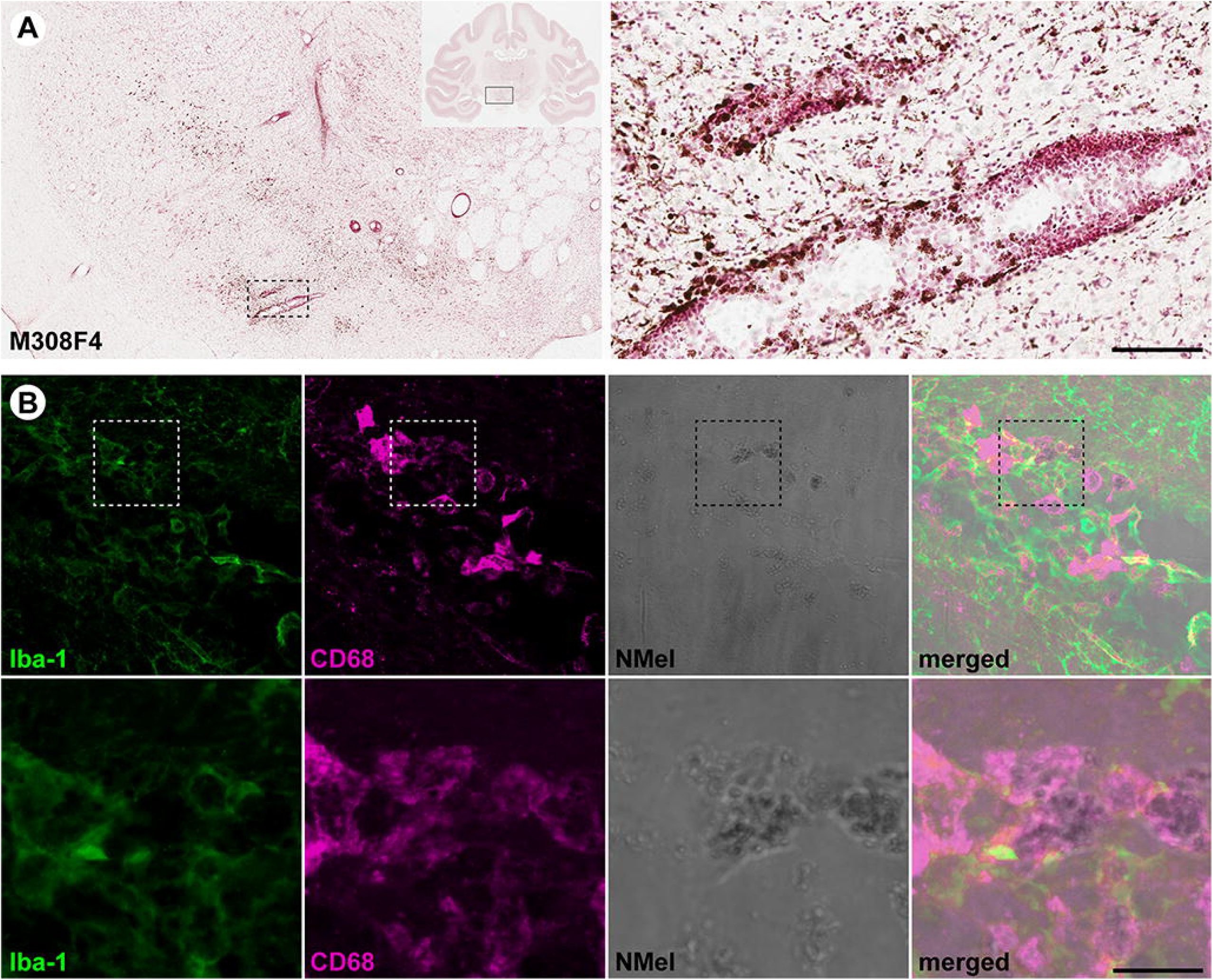
Extracellular neuromelanin accumulation and pro-inflammatory scenario. (A) Coronal section through the SNpc in animal M308F4 counterstained with neutral red and showing the distribution of extracellular NMel resulting from the loss of dopaminergic cells. Inset illustrates the preferential perivascular location of extracellular NMel deposits. Scale bar, 1000 μm (left panel) and 1000 μm (inset). (B) Confocal images showing the pro-inflammatory scenario made by cells positive for Iba-1 (green channel) and CD68 (red channel) that are digesting extracellular NMel. Scale bar, 64 μm (panels A-D) and 18 μm (panels A’-D’).

### Intracellular aggregates

Two different types of intracellular inclusions were constantly found in pigmented dopaminergic neurons, namely nuclear Marinesco bodies (MBs) and cytoplasmic Lewy bodies (LBs). Both types of inclusions are relatively abundant across the pigmented SNpc, and are easily identified by the immunofluorescent detection of P62, even at a low magnification (Figure 6A). Indeed, pigmented dopaminergic neurons showing between 1 to 3 inclusions per cell were often found. Although not all of NMel-containing TH+ neurons displayed intracellular inclusions, these structures were only observed in neuromelanized dopaminergic neurons. Accordingly, the presence of any potential relationship between intracellular NMel levels and the presence of intracellular inclusions (e.g. disclosing to what extent there was a pigmentation threshold needed to trigger the presence of inclusion bodies) was investigated under the confocal microscope at higher magnification, this analysis comprising randomly selected 25 pigmented neurons with inclusions and 25 pigmented neurons without P62+ intracellular aggregates in 12 consecutive sections of the SNpc covering the whole rostrocaudal extent of this structure. The conducted analysis revealed a lack of correlation between intracellular NMel densities and presence/absence of intracellular inclusions (Figure 6A). Although nuclear MBs and cytoplasmic LBs are both P62+ inclusions, only cytoplasmic LBs are also positive for α-Syn, meaning that besides exhibiting a different subcellular distribution, MBs and LBs can also be identified according to specific staining patterns (Figure 6B), as reported elsewhere^42,43^.

**Figure 6.**
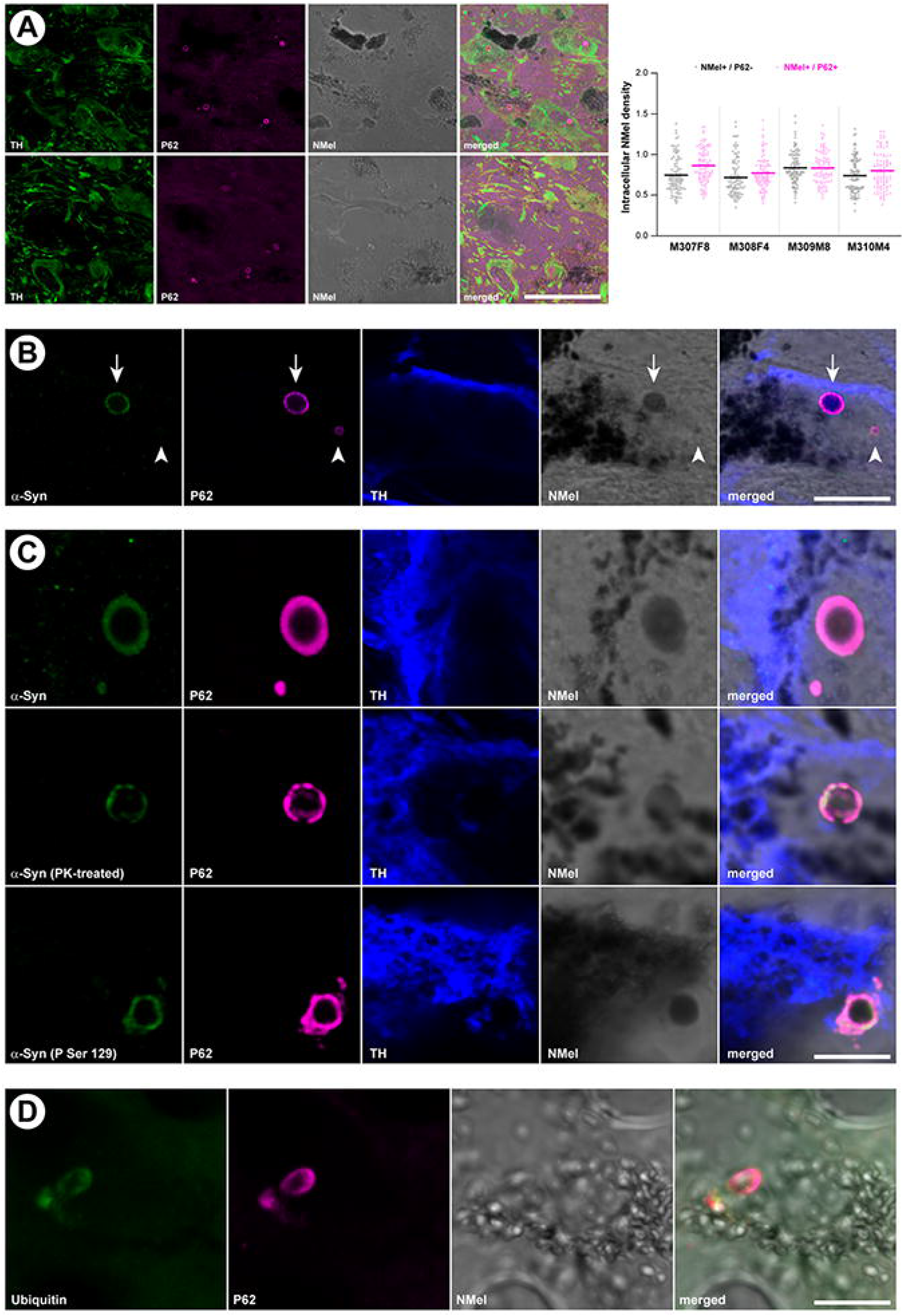
Intracellular inclusions in pigmented dopaminergic neurons of the substantia nigra. (A) Confocal images showing the relative abundance of intracellular inclusions within pigmented dopaminergic neurons in the SNpc. TH+ neurons are illustrated in the green channel and P62+ inclusions in the red channel. NMel is visualized under brightfield illumination. Images are taken from animal M310M4 (top) and M307F8 (bottom). Scale bar, 100 μm. The bloxplot illustrates NMel levels in pigmented neurons with and without P62+ intracellular inclusions. Analyses were conducted in 12 consecutive sections covering the entire rostrocaudal extent of the SNpc (equally spaced 400 μm each). (B) Two different types of intracellular inclusions were observed in pigmented dopaminergic neurons (TH+; blue channel), namely Marinesco bodies (MBs; intranuclear; arrowhead) and Lewy bodies (LBs; intracytoplasmic; arrow). While LBs are positive for both P62 (red channel) and α-Syn (green channel), MBs only displayed immunoreactivity for P62. Images are taken from animal M308F4. Scale bar, 15 μm. (C) LB-like intracytoplasmic inclusions within pigmented dopaminergic cells (TH+, blue channel) are made of insoluble and phosphorylated α-Syn species (green channel). Observed P62+ inclusions (red channel) are also labeled with antibodies against either total or phosphorylated α-Syn (top and bottom, respectively) and are resistant to digestion with proteinase K (middle row). All images are taken from animal M307F8. Scale bar, 5 μm. (D) Intracytoplasmic inclusions (e.g. LB-like) observed in neuromelanized neurons are positive for P62 (red channel) as well as for ubiquitin (green channel). Scale bar, 15 μm.

While the use of antibodies against total α-Syn revealed a complete co-localization between P62+ and α-Syn+ intracytoplasmic inclusions observed in pigmented dopaminergic neurons, to what extent these aggregates were made of insoluble forms of α-Syn was investigated by conducting either a pre-digestion of SNpc samples with proteinase K or by taking advantage of antibodies specifically detecting phosphorylated forms of α-Syn (P-Ser129). Both approaches revealed that P62+ LB cytoplasmic inclusions were made of pathological forms of α-Syn, either insoluble or phosphorylated (Figure 6C). Moreover, additional conducted stains also showed that P62+ inclusions are also positive for ubiquitin, another typical marker often used for identifying LBs (Figure 6D).

### Anterograde spread of endogenous alpha-synuclein

Upon having observed that the time-dependent NMel accumulation leads to the presence of intracytoplasmic inclusions made of endogenous α-Syn (LB-like), we sought to analyze the potential presence of similar inclusions in second-order pyramidal neurons of the prefrontal cortex receiving dopaminergic input. Intracytoplasmic inclusions positive for P62 and α-Syn were found in pyramidal neurons across the anterior cingulate gyrus, superior, middle and inferior frontal gyri and in the prefrontal gyrus of the left hemisphere. Within these cortical areas, P62+/ α-Syn+ inclusions were found in coronal section levels located between −2.0 and 6.0 mm of the anterior commissure, i.e. at a rostral distance from the injection sites in the SNpc ranging from 6.0 and 14.0 mm, respectively. Of particular importance, intracellular inclusions were only found in pyramidal neurons receiving TH+ fibers, often located in the post-synaptic element right opposite to the arrival of TH+ pre-synaptic boutons (Figure 7). Obtained results are providing experimental evidence showing that NMel-triggered endogenous synucleinopathy in midbrain dopaminergic neurons can spread in the anterograde direction through brain circuits, further inducing LB-like intracytoplasmic inclusions in second-order cortical pyramidal neurons through permissive trans-synaptic templating.

**Figure 7.**
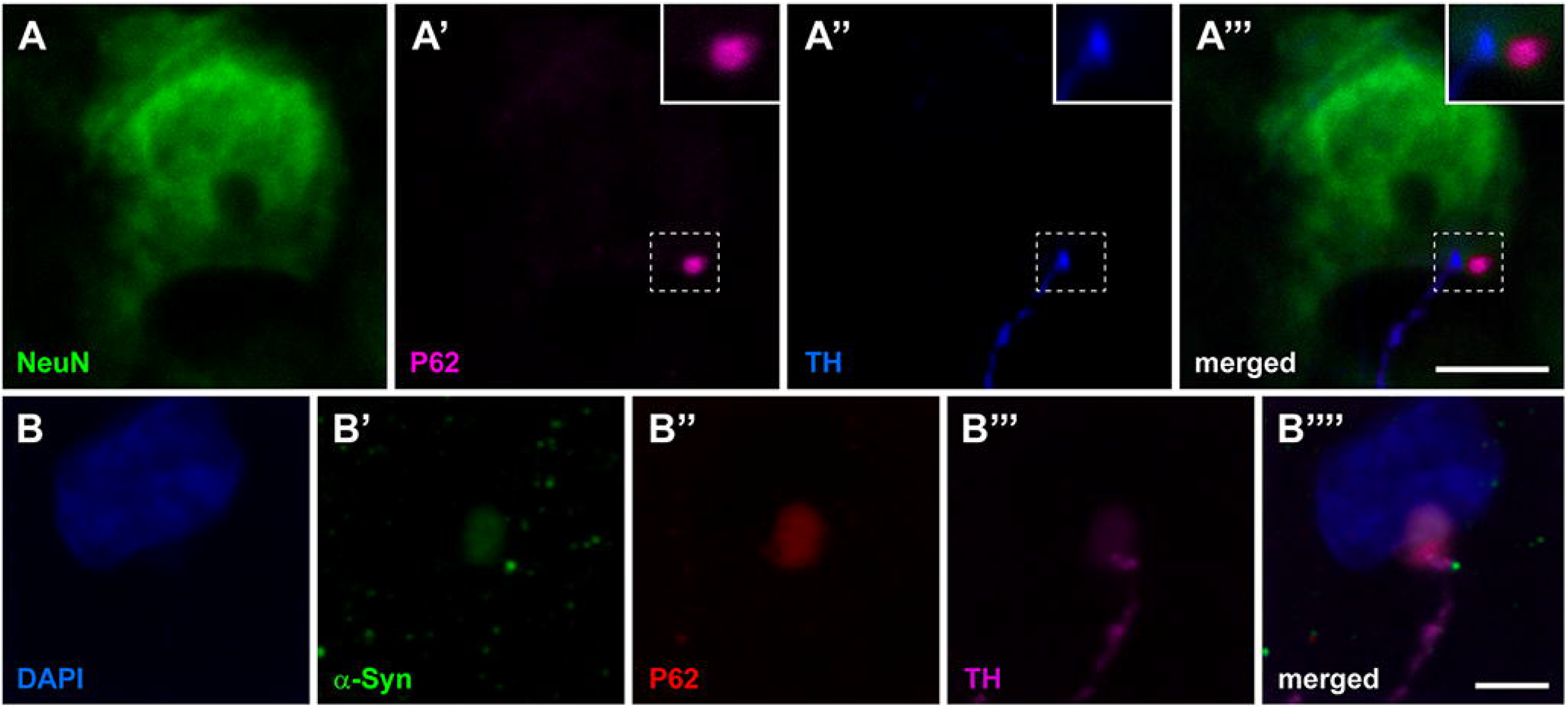
Anterograde spread of endogenous alpha-synuclein towards the prefrontal cerebral cortex. (A-A’’’) Confocal images taken at the level of the superior frontal gyrus in animal M309M8 showing a layer V pyramidal neuron (stained with NeuN; green channel) receiving a TH+ terminal (blue channel) and showing an intracytoplasmic inclusion (positive for P62) just opposite to the arrival of the terminal bouton. Scale bar, 10 μm. (B-B’’’) Confocal images taken from the anterior cingulate gyrus in animal M307F8 showing that P62+ intracytoplasmic inclusions (red channel) are also positive for α-Syn (green channel) and are observed in layer V neurons receiving TH+ terminals (purple channel). Scale bar, 20 μm.

## Discussion

Here a novel NHP animal model of Parkinson’s disease has been developed and characterized, this model mimicking the known neuropathological hallmarks of the disorder with unprecedented accuracy. The AAV-mediated enhanced expression of human tyrosinase gene induced a pigmentation of dopaminergic neurons in the SNpc up to similar levels observed in elderly humans. Next, a time-dependent loss of pigmentation was observed, resulting from progressive degeneration of nigrostriatal neurons. Furthermore, intracellular inclusions (Marinesco and Lewy bodies) were found within NMel-containing dopaminergic cells, together with a pro-inflammatory scenario mediated by microglial cells and macrophages. While the main aim of the conducted study was to generate a novel NHP model of PD, mechanistic evidence showing that the enhanced NMel accumulation triggered an endogenous synucleinopathy in the SNpc is also provided. Moreover, obtained data supported the experimental demonstration of the Braak hypothesis, by showing the presence of an anterograde spread of the endogenous synucleinopathy towards neurons of the prefrontal cortex receiving dopaminergic input through permissive trans-synaptic templating.

### Technical considerations and limitations of the study

The current worldwide shortage of NHP supplies^45^ forced us to use a small number of experimental subjects, therefore several questions remained open, at least to some extent. For instance, to what extent direct stereotaxic deliveries of AAVs in the SNpc may have induced any inherent damage in the targeted structure leading to neuronal death and gliosis cannot be ruled out. Indeed, conducted microPET neuroimage studies revealed a time-dependent reduction of radiotracer binding potential in both hemispheres of female animals when compared to baseline levels (however the reduction was always more prominent in the putamen nucleus in the left hemisphere). Most importantly -and unexpectedly-loss of dopaminergic cells and nigrostriatal terminals was only noticed in female animals at 4 and 8 months post-delivery of AAV-hTyr (animals M308F4 and M307F8, respectively), therefore raising the question of to what extent there may be a gender effect. Regarding male subjects, a mild cell loss in the SNpc (19.93%) was only observed after a follow-up period of 8 months (animal M308M8), whereas an almost negligible 8.63% of dopaminergic damage was found 4 months post-injection of AAV-hTyr (animal M310M4). Such small percentages of dopaminergic cell degeneration lead to a non-significant reduction of nigrostriatal terminals in the caudate and putamen nuclei in these animals. Furthermore, it is also worth noting that female animals exhibited higher mean intracellular levels of NMel than male subjects. Any other outcomes of the conducted procedure, such as percentages of transduced dopaminergic neurons with AAV-hTyr, intracellular inclusions, gliosis and anterograde spread of α-Syn towards the prefrontal cortex, were always observed in all animals. In summary, the question arising here is why male subjects, showing similar neuropathological signatures than females, seem to be more resistant to dopaminergic neurodegeneration. Our initial thought was that different accuracies for the stereotaxic delivery of AAV-hTyr across animals might account for the observed phenomena, bearing in mind that the SNpc is a demanding stereotaxic target, deeply located in the ventral mesencephalon. However, a careful analysis of the injection sites (2 sites in the left SNpc for AAV-hTyr injections and two sites in the right SNpc for deliveries of the control vector) revealed that performed stereotaxic deposits were equivalent across animals, with a minimal-if any-inter-individual variability. Moreover, gender differences were not reported in available AAV-mediated NHP models of synucleinopathy, such as those conducted in marmosets made of an equal number of male and female subjects^46,47^. Similar approaches performed in *M. fascicularis*^28^, only tested AAV2-SynA53T deliveries in a cohort of female macaques, whereas a more recent study conducted in our group with AAV9-SynA53T^32^ only comprised male animals, and therefore the presence of a gender effect cannot be assessed. Finally, a different study analyzing the patterns of age-dependent physiological pigmentation with NMel in the human SNpc did not reported any gender effect in individuals spanning the ages of 24 weeks to 95 years old^2^.

While this model was designed to reproduce the neuropathological scenario of human PD to the best possible extent, another limitation is represented by the lack of any noticeable motor phenotype. The development of a motor syndrome was not expected with percentages of dopaminergic cell loss barely reaching 50% in females and less than 20% in male animals. Indeed, NHPs are very efficient in self-compensating unilateral insults. According to our former, long-time experience in the NHP model with MPTP, a bilateral damage of nigrostriatal-projecting dopaminergic neurons above 85% is needed to induce a motor phenotype^48,49^. Studies performed in marmosets with AAVs coding for either wild-type or mutated forms of α-Syn^46,47^ revealed that a motor phenotype cannot be appreciated before 9 weeks post-inoculation of the viral suspension, later leading to head position bias and body rotations between 15 and 27 weeks. Furthermore, motor phenotypes were not reported upon the delivery of preformed α-Syn fibrils in the putamen and caudate nuclei of two female marmosets 12 weeks post-injection^29^. The same holds true for experiments conducted in *M. fascicularis* with the AAV-mediated enhancement of α-Syn in the SNpc with follow-up periods ranging between 12 and 17 weeks^27,28,32^.

### Available NHP models of synucleinopathy

Although broadly used as the gold-standard choice, traditional neurotoxin-based models of PD in NHPs failed to recapitulate the main neuropathological events that typically characterize PD^50^. In an attempt to get closer to the human condition, the field has moved towards the use AAVs coding for either wild-type or mutated forms of α-Syn in NHPs^30^. When conducted this way, a variable 30-60% loss of dopaminergic cells together with a well-established α-Syn pathology was observed in marmosets following 16 weeks post-injection of AAV2-SynWT^46^. Next, the use of a different AAV serotype (AAV2/5 instead of AAV2) in marmosets lead a constant cell loss above 40%^47^. Lower levels of dopaminergic cell loss were reported in marmosets 11 weeks after the delivery of AAV9-SynA53T, reaching 13% in young animals and 20% in older animals^27^. A similar approach conducted in macaques (*M. fascicularis*), with intraparenchymal deliveries of AAV2 coding for mutated α-Syn in the SNpc lead to a 50% dopaminergic cell loss after a follow-up period of 17 weeks post-injection^28^. Former data from our group comprising the delivery of AAV9-SynA53T into the SNpc of *M.fascicularis*, reported a 39% of cell loss in the SNpc 12 weeks post-injection of the viral vector^32^. Moreover, a different approach for modeling synucleinopathy in NHPs has been made available by the intrastriatal delivery of synthetic human α-Syn fibrils^29,31^. In two female marmosets, this strategy lead to PD-like α-Syn pathology 12 weeks post-injection together with non-quantified levels of dopaminergic cell death^29^. Regarding the study conducted in one male *M. fuscata*^31^, a mild neuronal cell loss in the SNpc together with the presence of LB-like intracytoplasmic inclusions and gliosis 12 weeks post-delivery of α-Syn fibrils were observed, these pathologies being more evident in the peri-injected areas of the putamen nucleus. Compared to strategies using AAVs coding for α-Syn, data reported here showed similar ranges of dopaminergic cell loss in female animals with a follow-up of either 4 or 8 months post-injection of AAV-hTyr (animals M308F4 and M307F8, respectively). However, lower levels of dopaminergic neurodegeneration were observed in both male animals, reaching up to 19.93% of loss after 8 months post-delivery of the viral vector (animal M309M8), and only 8.63% with a follow-up of 4 months (animal M310M4).

### Accuracy of the model in mimicking the known neuropathology of human PD

Although at a more costly option, setting up better NHP models of PD remain fundamental for pushing forward disease-modifying therapeutics in an attempt to minimize clinical trial failures as well as in gaining more insight on disease biology^45,51^. In this regard, an ideal NHP model should reproduce all neuropathological signatures that typically account for human PD, such as a progressive loss of pigmented dopaminergic cells in the SNpc and intracellular inclusions. Moreover, although primates (human and non-human) are the only animal species showing a time-dependent macroscopic pigmentation in the SNpc^6^, such a progressive pigmentation is physiological, with light NMel granular deposits being detectable at approximately 3 years of age in humans^2^. In NHPs, a mild pigmentation can only be detected with optical microscopy in squirrel monkeys (*Saimiri scireus*) at 6 years of age^52^, and at 8 years of age in Rhesus macaques (MacBrain Resource Collections; https://macbraingallery.yale.edu/). Pigmentation gradually increases with age, particularly in humans^2^ and to a lesser extent in macaques (MacBrain Resource Collections; https://macbraingallery.yale.edu/), although age-related changes have not been observed in squirrel monkeys between 6, 12 and 21 years^52^. Regarding juvenile long-tailed macaques (*M. fascicularis*) as the ones used here (approximately 4 years-old), NMel pigmentation is not expected, even at the microscopic level. In these animals, the delivery of AAV-hTyr into the SNpc, lead to macro- and microscopic pigmentation levels similar to elderly humans. However, neuropathological signatures reported here resulting from the forced expression of human tyrosinase gene cannot be taken as physiological. We hypothesize that the abnormal NMel accumulation is leading to an accelerated aged phenotype. Aging is the main risk factor in developing PD and it has long been known that dopaminergic neurons are more susceptible to senescence than any other neuronal types^53–57^ and indeed aged macaques are more sensitive to MPTP treatment^58–60^. Moreover, an age-dependent marked increase in α-Syn content within dopaminergic cells was reported in rhesus macaques and humans^61^. Another observation pointing into this direction is the relative abundance of Marinesco bodies (MBs) found here in pigmented dopaminergic neurons. Often considered as an indicator of advanced age^62^, MBs are more frequently observed in nigral neurons^63,64^, in particular within those pigmented neurons also exhibiting LBs, a finding suggesting a pathological role for MBs^22^ contrary to earlier views considering MBs as age-related incidental findings^65^. Out data are in keeping with available evidence, by showing that MBs are present in pigmented neurons with LBs and are not stained with antibodies against α-Syn^42,43^.

### Anterograde spread of endogenous alpha-synuclein

While not designed for this purpose, obtained data support the fact that the progressive NMel accumulation in dopaminergic neurons triggers an endogenous synucleinopathy. Importantly, such an endogenous synucleinopathy is able to progress anterogradely towards pyramidal neurons of the prefrontal cortex receiving dopaminergic input, therefore suggesting a potential neuron-to-neuron circuit-specific spread of α-Syn through permissive trans-synaptic templating, in keeping with the so-called Braak hypothesis^24,25,66^. Although there is a close correlation between neuropathological staging and clinical disease course, to what extent the Braak hypothesis is either pathophysiologically relevant or an epiphenomenon within the context of PD remains controversial^67–69^. Central to this controversy is the lack of experimental evidence for a prionoid-like non-random spread of endogenous α-Syn. Initial evidence arising from clinical trials in PD patients receiving mesencephalic cell grafts^70,71^ was later on reproduced in mice and macaques receiving brain extracts from PD patients^72^. Although there are convincing experimental data both at the *in vivo* and *in vitro* levels in support of the prionoid spread of α-Syn^73–76^, available evidence is based on the administration of exogenous α-Syn species (e.g. either AAVs encoding the SNCA gene, brain extracts of PD patients or preformed α-Syn fibrils), and therefore to what extent α-Syn pathology of endogenous origins can spread remains to be proved^69^. Without the aim of adding more controversy to an already heated debate, data obtained here provided support for spreading phenomena of endogenous synucleinopathy-triggered by NMel accumulation-from pigmented dopaminergic cells towards pyramidal neurons of the prefrontal cortex in NHPs.

### Conclusions

The pigmented NHP model introduced here mimics the known neuropathological hallmarks of human PD with unprecedented accuracy. Moreover, obtained evidence showed that the time-dependent NMel accumulation triggers the pathological misfolding of endogenous α-Syn in pigmented midbrain dopaminergic cells and indeed such synucleinopathy is able to spread anterogradely towards the cerebral cortex. Recent failures in clinical trials targeting α-Syn^77,78^ called for a reappraisal of the underlying rationale^79^. Bearing in mind that NMel accumulation seems to be responsible for α-Syn pathology, it is perhaps worth considering alternative approaches intended to reduce intracellular NMel levels in an attempt to get rid of the subsequent synucleinopathy. Supporting this concept, reduction of intracellular NMel levels in vivo, either by boosting NMel cytosolic clearance with the autophagy activator TFEB^34^ or by reducing NMel production with VMAT2-mediated enhancement of dopamine vesicular encapsulation^80^, resulted in a major attenuation of the PD phenotype, both at the behavioral and neuropathological levels, in AAV-hTyr-injected NMel-producing rats. Therefore, strategies decreasing age-dependent NMel building up may provide unprecedented therapeutic opportunities to either prevent, halt or delay neuronal dysfunction and degeneration linked to PD and, in a broader sense, brain aging.

## Supporting information

plasmid map (pAAV-CMV-hTyr)

sequence for pAAV-CMV-hTyr

Animal records

raw data Figure 2

raw data Figure 2

raw data Figure 2

raw data Figure 2

raw data Figure 2

raw data Figure 3C2

raw data Figure 3C3

raw data Figue 4A

raw data figure 4B

raw data figure 6A

## Abbreviations

AAVs: adeno-associated viral vectors
ac: anterior commissure
ac-pc plane: bicommissural plane
α-Syn: alpha-synuclein
CD68: cluster of differentiation 68
CMV: cytomegalovirus
DMSO: dimethylsulphoxide
^11^C-DTBZ: dihydrotetrabenazine
hTyr: human tyrosinase gene
Iba-1: ionized calcium-binding adapter molecule 1
ITRs: inverted terminal repeats
LBs: Lewy bodies
NHP: non-human primates
NMel: neuromelanin
MBs: Marinesco bodies
MPTP: 1-methyl-4-phenyl-1,2,3,6-tetrahydropyridine
NeuN: neuronal marker
NR: neutral red
PB: phosphate buffer
PD: Parkinson’s disease
ROIs: regions of interest
SNCA: alpha-synuclein gene
SNpc: Substantia nigra pars compacta
TFEB: transcription factor EB
TH: tyrosine hydroxylase
VMAT2: vesicular monoamine transporter type 2.

## Acknowledgements

The authors acknowledge the support received from Marta González Sepúlveda, postdoctoral fellow at Vall d’Hebron Research Institute, Autonomous University of Barcelona, Spain.

## Supplementary material

Supplementary material is available at *Brain* online.

---

Supplementary Figure 1. Plasmid map for pAAV-CMV-hTyr

Supplementary Figure 2. Sequence for pAAV-CMV-hTyr

