## Supplementary figures and images for "Neuromelanin accumulation drives endogenous synucleinopathy in non-human primates"

### plasmid map (pAAV-CMV-hTyr)

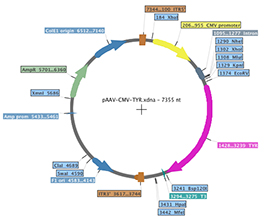
