## Supplementary material for "Neuromelanin accumulation drives endogenous synucleinopathy in non-human primates": sequence for pAAV-CMV-hTyr

pAAV-CMV-Tyr Sequence

CAGCAGCTGCGCGCTCGCTCGCTCACTGAGGCCGCCCGGGCAAAGCCCGGGCGTCGGGCGACCTTTGGTCGCCCGGCCTC

AGTGAGCGAGCGAGCGCGCAGAGAGGGAGTGGCCAACTCCATCACTAGGGGTTCCTTGTAGTTAATGATTAACCCGCCAT

GCTACTTATCTACGTAGCCATGCTCTAGACATGGCTCGACAGATCTCAATATTGGCCATTAGCCATATTATTCATTGGTT

ATATAGCATAAATCAATATTGGCTATTGGCCATTGCATACGTTGTATCTATATCATAATATGTACATTTATATTGGCTCA

TGTCCAATATGACCGCCATGTTGGCATTGATTATTGACTAGTTATTAATAGTAATCAATTACGGGGTCATTAGTTCATAG

CCCATATATGGAGTTCCGCGTTACATAACTTACGGTAAATGGCCCGCCTGGCTGACCGCCCAACGACCCCCGCCCATTGA

CGTCAATAATGACGTATGTTCCCATAGTAACGCCAATAGGGACTTTCCATTGACGTCAATGGGTGGAGTATTTACGGTAA

ACTGCCCACTTGGCAGTACATCAAGTGTATCATATGCCAAGTCCGCCCCCTATTGACGTCAATGACGGTAAATGGCCCGC

CTGGCATTATGCCCAGTACATGACCTTACGGGACTTTCCTACTTGGCAGTACATCTACGTATTAGTCATCGCTATTACCA

TGGTGATGCGGTTTTGGCAGTACACCAATGGGCGTGGATAGCGGTTTGACTCACGGGGATTTCCAAGTCTCCACCCCATT

GACGTCAATGGGAGTTTGTTTTGGCACCAAAATCAACGGGACTTTCCAAAATGTCGTAACAACTGCGATCGCCCGCCCCG

TTGACGCAAATGGGCGGTAGGCGTGTACGGTGGGAGGTCTATATAAGCAGAGCTCGTTTAGTGAACCGTCAGATCACTAG

AAGCTTTATTGCGGTAGTTTATCACAGTTAAATTGCTAACGCAGTCAGTGCTTCTGACACAACAGTCTCGAACTTAAGCT

GCAGTGACTCTCTTAAGGTAGCCTTGCAGAAGTTGGTCGTGAGGCACTGGGCAGGTAAGTATCAAGGTTACAAGACAGGT

TTAAGGAGACCAATAGAAACTGGGCTTGTCGAGACAGAGAAGACTCTTGCGTTTCTGATAGGCACCTATTGGTCTTACTG

ACATCCACTTTGCCTTTCTCTCCACAGGTGTCCACTCCCAGTTCAATTACAGCTCTTAAGGCTAGAGTACTTAATACGAC

TCACTATAGGCTAGCctcgacctcgagacgcgtgataaacttaagcttggtaccgagctcggatccactagtccagtgtg

gtggaattctgcagatatccagcacagtggcggccgctcgaccGACCTTGTGAGGACTAGAGGAAGAATGCTCCTGGCTG

TTTTGTACTGCCTGCTGTGGAGTTTCCAGACCTCCGCTGGCCATTTCCCTAGAGCCTGTGTCTCCTCTAAGAACCTGATG

GAGAAGGAATGCTGTCCACCGTGGAGCGGGGACAGGAGTCCCTGTGGCCAGCTTTCAGGCAGAGGTTCCTGTCAGAATAT

CCTTCTGTCCAATGCACCACTTGGGCCTCAATTTCCCTTCACAGGGGTGGATGACCGGGAGTCGTGGCCTTCCGTCTTTT

ATAATAGGACCTGCCAGTGCTCTGGCAACTTCATGGGATTCAACTGTGGAAACTGCAAGTTTGGCTTTTGGGGACCAAAC

TGCACAGAGAGACGACTCTTGGTGAGAAGAAACATCTTCGATTTGAGTGCCCCAGAGAAGGACAAATTTTTTGCCTACCT

CACTTTAGCAAAGCATACCATCAGCTCAGACTATGTCATCCCCATAGGGACCTATGGCCAAATGAAAAATGGATCAACAC

CCATGTTTAACGACATCAATATTTATGACCTCTTTGTCTGGATGCATTATTATGTGTCAATGGATGCACTGCTTGGGGGA

TCTGAAATCTGGAGAGACATTGATTTTGCCCATGAAGCACCAGCTTTTCTGCCTTGGCATAGACTCTTCTTGTTGCGGTG

GGAACAAGAAATCCAGAAGCTGACAGGAGATGAAAACTTCACTATTCCATATTGGGACTGGCGGGATGCAGAAAAGTGTG

ACATTTGCACAGATGAGTACATGGGAGGTCAGCACCCCACAAATCCTAACTTACTCAGCCCAGCATCATTCTTCTCCTCT

TGGCAGATTGTCTGTAGCCGATTGGAGGAGTACAACAGCCATCAGTCTTTATGCAATGGAACGCCCGAGGGACCTTTACG

GCGTAATCCTGGAAACCATGACAAATCCAGAACCCCAAGGCTCCCCTCTTCAGCTGATGTAGAATTTTGCCTGAGTTTGA

CCCAATATGAATCTGGTTCCATGGATAAAGCTGCCAATTTCAGCTTTAGAAATACACTGGAAGGATTTGCTAGTCCACTT

ACTGGGATAGCGGATGCCTCTCAAAGCAGCATGCACAATGCCTTGCACATCTATATGAATGGAACAATGTCCCAGGTACA

GGGATCTGCCAACGATCCTATCTTCCTTCTTCACCATGCATTTGTTGACAGTATTTTTGAGCAGTGGCTCCGAAGGCACC

GTCCTCTTCAAGAAGTTTATCCAGAAGCCAATGCACCCATTGGACATAACCGGGAATCCTACATGGTTCCTTTTATACCA

CTGTACAGAAATGGTGATTTCTTTATTTCATCCAAAGATCTGGGCTATGACTATAGCTATCTACAAGATTCAGACCCAGA

CTCTTTTCAAGACTACATTAAGTCCTATTTGGAACAAGCGAGTCGGATCTGGTCATGGCTCCTTGGGGCGGCGATGGTAG

GGGCCGTCCTCACTGCCCTGCTGGCAGGGCTTGTGAGCTTGCTGTGTCGTCACAAGAGAAAGCAGCTTCCTGAAGAAAAG

CAGCCACTCCTCATGGAGAAAGAGGATTACCACAGCTTGTATCAGAGCCATTTATAAAAGGCTTAGGCAATAGAGTAGGG

CCAAAAAGCCTGACCTCACTCTAACTCAAAGTAATGTCCAGGTTCCCAGAGAATATCTGCTGGTATTTTTCTGTAAAGAC

CATTTGCAAAATTGTAACCTAATACAAAGTGTAGCCTTCTTCCAACTCAGGTAGAACACACCTGTCTTTGTCTTGCTGTT

TTCACTCAGCCCTTTTAACATTTTCCCCTAAGCCCctagagggcccgtttatcggatcccggccggcggccgcTTCCCTT

TAGTGAGGGTTAATGCTTCGAGCAGACATGATAAGATACATTGATGAGTTTGGACAAACCACAACTAGAATGCAGTGAAA

AAAATGCTTTATTTGTGAAATTTGTGATGCTATTGCTTTATTTGTAACCATTATAAGCTGCAATAAACAAGTTAACAACA

ACAATTGCATTCATTTTATGTTTCAGGTTCAGGGGGAGATGTGGGAGGTTTTTTAAAGCAAGTAAAACCTCTACAAATGT

GGTAAAATCCGATAAGGGACTAGAGCATGGCTACGTAGATAAGTAGCATGGCGGGTTAATCATTAACTACAAGGAACCCC

TAGTGATGGAGTTGGCCACTCCCTCTCTGCGCGCTCGCTCGCTCACTGAGGCCGGGCGACCAAAGGTCGCCCGACGCCCG

GGCTTTGCCCGGGCGGCCTCAGTGAGCGAGCGAGCGCGCCAGCTGGCGTAATAGCGAAGAGGCCCGCACCGATCGCCCTT

CCCAACAGTTGCGCAGCCTGAATGGCGAATGGAATTCCAGACGATTGAGCGTCAAAATGTAGGTATTTCCATGAGCGTTT

TTCCGTTGCAATGGCTGGCGGTAATATTGTTCTGGATATTACCAGCAAGGCCGATAGTTTGAGTTCTTCTACTCAGGCAA

GTGATGTTATTACTAATCAAAGAAGTATTGCGACAACGGTTAATTTGCGTGATGGACAGACTCTTTTACTCGGTGGCCTC

ACTGATTATAAAAACACTTCTCAGGATTCTGGCGTACCGTTCCTGTCTAAAATCCCTTTAATCGGCCTCCTGTTTAGCTC

CCGCTCTGATTCTAACGAGGAAAGCACGTTATACGTGCTCGTCAAAGCAACCATAGTACGCGCCCTGTAGCGGCGCATTA

AGCGCGGCGGGTGTGGTGGTTACGCGCAGCGTGACCGCTACACTTGCCAGCGCCCTAGCGCCCGCTCCTTTCGCTTTCTT

CCCTTCCTTTCTCGCCACGTTCGCCGGCTTTCCCCGTCAAGCTCTAAATCGGGGGCTCCCTTTAGGGTTCCGATTTAGTG

CTTTACGGCACCTCGACCCCAAAAAACTTGATTAGGGTGATGGTTCACGTAGTGGGCCATCGCCCTGATAGACGGTTTTT

CGCCCTTTGACGTTGGAGTCCACGTTCTTTAATAGTGGACTCTTGTTCCAAACTGGAACAACACTCAACCCTATCTCGGT

CTATTCTTTTGATTTATAAGGGATTTTGCCGATTTCGGCCTATTGGTTAAAAAATGAGCTGATTTAACAAAAATTTAACG

CGAATTTTAACAAAATATTAACGTCTACAATTTAAATATTTGCTTATACAATCTTCCTGTTTTTGGGGCTTTTCTGATTA

TCAACCGGGGTACATATGATTGACATGCTAGTTTTACGATTACCGTTCATCGATTCTCTTGTTTGCTCCAGACTCTCAGG

CAATGACCTGATAGCCTTTGTAGAGACCTCTCAAAAATAGCTACCCTCTCCGGCATGAATTTATCAGCTAGAACGGTTGA

ATATCATATTGATGGTGATTTGACTGTCTCCGGCCTTTCTCACCCGTTTGAATCTTTACCTACACATTACTCAGGCATTG

CATTTAAAATATATGAGGGTTCTAAAAATTTTTATCCTTGCGTTGAAATAAAGGCTTCTCCCGCAAAAGTATTACAGGGT

CATAATGTTTTTGGTACAACCGATTTAGCTTTATGCTCTGAGGCTTTATTGCTTAATTTTGCTAATTCTTTGCCTTGCCT

GTATGATTTATTGGATGTTGGAATCGCCTGATGCGGTATTTTCTCCTTACGCATCTGTGCGGTATTTCACACCGCATATG

GTGCACTCTCAGTACAATCTGCTCTGATGCCGCATAGTTAAGCCAGCCCCGACACCCGCCAACACCCGCTGACGCGCCCT

GACGGGCTTGTCTGCTCCCGGCATCCGCTTACAGACAAGCTGTGACCGTCTCCGGGAGCTGCATGTGTCAGAGGTTTTCA

CCGTCATCACCGAAACGCGCGAGACGAAAGGGCCTCGTGATACGCCTATTTTTATAGGTTAATGTCATGATAATAATGGT

TTCTTAGACGTCAGGTGGCACTTTTCGGGGAAATGTGCGCGGAACCCCTATTTGTTTATTTTTCTAAATACATTCAAATA

TGTATCCGCTCATGAGACAATAACCCTGATAAATGCTTCAATAATATTGAAAAAGGAAGAGTATGAGTATTCAACATTTC

CGTGTCGCCCTTATTCCCTTTTTTGCGGCATTTTGCCTTCCTGTTTTTGCTCACCCAGAAACGCTGGTGAAAGTAAAAGA

TGCTGAAGATCAGTTGGGTGCACGAGTGGGTTACATCGAACTGGATCTCAACAGCGGTAAGATCCTTGAGAGTTTTCGCC

CCGAAGAACGTTTTCCAATGATGAGCACTTTTAAAGTTCTGCTATGTGGCGCGGTATTATCCCGTATTGACGCCGGGCAA

GAGCAACTCGGTCGCCGCATACACTATTCTCAGAATGACTTGGTTGAGTACTCACCAGTCACAGAAAAGCATCTTACGGA

TGGCATGACAGTAAGAGAATTATGCAGTGCTGCCATAACCATGAGTGATAACACTGCGGCCAACTTACTTCTGACAACGA

TCGGAGGACCGAAGGAGCTAACCGCTTTTTTGCACAACATGGGGGATCATGTAACTCGCCTTGATCGTTGGGAACCGGAG

CTGAATGAAGCCATACCAAACGACGAGCGTGACACCACGATGCCTGTAGCAATGGCAACAACGTTGCGCAAACTATTAAC

TGGCGAACTACTTACTCTAGCTTCCCGGCAACAATTAATAGACTGGATGGAGGCGGATAAAGTTGCAGGACCACTTCTGC

GCTCGGCCCTTCCGGCTGGCTGGTTTATTGCTGATAAATCTGGAGCCGGTGAGCGTGGGTCTCGCGGTATCATTGCAGCA

CTGGGGCCAGATGGTAAGCCCTCCCGTATCGTAGTTATCTACACGACGGGGAGTCAGGCAACTATGGATGAACGAAATAG

ACAGATCGCTGAGATAGGTGCCTCACTGATTAAGCATTGGTAACTGTCAGACCAAGTTTACTCATATATACTTTAGATTG

ATTTAAAACTTCATTTTTAATTTAAAAGGATCTAGGTGAAGATCCTTTTTGATAATCTCATGACCAAAATCCCTTAACGT

GAGTTTTCGTTCCACTGAGCGTCAGACCCCGTAGAAAAGATCAAAGGATCTTCTTGAGATCCTTTTTTTCTGCGCGTAAT

CTGCTGCTTGCAAACAAAAAAACCACCGCTACCAGCGGTGGTTTGTTTGCCGGATCAAGAGCTACCAACTCTTTTTCCGA

AGGTAACTGGCTTCAGCAGAGCGCAGATACCAAATACTGTCCTTCTAGTGTAGCCGTAGTTAGGCCACCACTTCAAGAAC

TCTGTAGCACCGCCTACATACCTCGCTCTGCTAATCCTGTTACCAGTGGCTGCTGCCAGTGGCGATAAGTCGTGTCTTAC

CGGGTTGGACTCAAGACGATAGTTACCGGATAAGGCGCAGCGGTCGGGCTGAACGGGGGGTTCGTGCACACAGCCCAGCT

TGGAGCGAACGACCTACACCGAACTGAGATACCTACAGCGTGAGCTATGAGAAAGCGCCACGCTTCCCGAAGGGAGAAAG

GCGGACAGGTATCCGGTAAGCGGCAGGGTCGGAACAGGAGAGCGCACGAGGGAGCTTCCAGGGGGAAACGCCTGGTATCT

TTATAGTCCTGTCGGGTTTCGCCACCTCTGACTTGAGCGTCGATTTTTGTGATGCTCGTCAGGGGGGCGGAGCCTATGGA

AAAACGCCAGCAACGCGGCCTTTTTACGGTTCCTGGCCTTTTGCTGGCCTTTTGCTCACATGTTCTTTCCTGCGTTATCC

CCTGATTCTGTGGATAACCGTATTACCGCCTTTGAGTGAGCTGATACCGCTCGCCGCAGCCGAACGACCGAGCGCAGCGA

GTCAGTGAGCGAGGAAGCGGAAGAGCGCCCAATACGCAAACCGCCTCTCCCCGCGCGTTGGCCGATTCATTAATG
